# Air pollution drives macrophage senescence through a phagolysosome-15-lipoxygenase pathway

**DOI:** 10.1101/2024.01.04.574228

**Authors:** Sarah A. Thomas, Hwan Mee Yong, Ana M. Rule, Naina Gour, Stephane Lajoie

## Abstract

Urban particulate matter (uPM) poses significant health risks, particularly to the respiratory system. Fine particles, such as PM_2.5_, can penetrate deep into the lungs and exacerbate a range of health problems, including emphysema, asthma, and lung cancer. PM exposure is also linked to extra-pulmonary disorders like heart and neurodegenerative diseases. Moreover, prolonged exposure to elevated PM levels can reduce overall life expectancy. Senescence is a dysfunctional cell state typically associated with age but can also be precipitated by environmental stressors. This study aimed to determine whether uPM could drive senescence in macrophages, an essential cell type involved in particulate phagocytosis-mediated clearance. While it is known that uPM exposure impairs immune function, this deficit is multi-faceted and incompletely understood, partly due to the use of particulates such as diesel exhaust particle (DEP) as a surrogate for true uPM. uPM was collected from several locations in the USA, including Baltimore, Houston, and Phoenix. Bone marrow-derived macrophages (BMDMs) were stimulated with uPM or reference particulates (*e.g.,* DEP) to assess senescence-related parameters. We report that uPM-exposed BMDMs adopt a senescent phenotype characterized by increased IL-1α secretion, senescence-associated β-galactosidase activity, and diminished proliferation. Exposure to allergens failed to elicit such a response, supporting a distinction between different types of environmental exposures. uPM-induced senescence was independent of key macrophage activation pathways, specifically inflammasome and scavenger receptor. However, inhibition of the phagolysosome pathway abrogated senescence markers, supporting this phenotype’s attribution to uPM phagocytosis. These data suggest uPM exposure leads to macrophage senescence, which may contribute to immunopathology.

## Introduction

Urban particulate matter (uPM) pollution significantly threatens human health. The American Lung Association’s annual State of the Air 2023 report suggests that one-third of Americans live in areas with a failing grade for air pollution (*State of the Air*, 2023). Globally, it has been estimated that fossil-fuel-derived PM_2.5_ is responsible for as many as 10 million deaths annually (Vohra et al., 2021), and increasing air pollution is estimated to affect half of the world’s population (Shaddick et al., 2020). Fossil fuels contribute to PM directly and indirectly through climate change’s ecological effects. In particular, the contribution of wildfire smoke to PM_2.5_ levels has recently been highlighted as a key obstacle to improving air quality in the US (Burke et al., 2023). The association between PM exposure and poor health outcomes is well-established, with multi-system implications including respiratory, neurological, and cardiac sequelae (*State of the Air*, 2023; Thangavel et al., 2022). Notably, even acute uPM exposure is sufficient to cause harm and has been associated with an increased risk of myocardial infarction (Bhaskaran et al., 2011).

While uPM exposure is associated with immune dysfunction (Adami et al., 2022; Li et al., 2022; Ural et al., 2022), the molecular mechanisms driving such dysfunction are incompletely understood. Macrophages are highly phagocytic immune cells and are the primary cells responsible for responding to environmental exposures (Ross et al., 2021). PM exposure has been linked to senescence in various cell types, including fibroblasts (Sachdeva et al., 2019); this association has yet to be explored in the context of macrophages. Senescence is usually considered in the context of aging but can also be precipitated by environmental stressors, known as disease-related senescence (Childs et al., 2015). Given the sub-optimal functionality of senescent cells, which no longer undergo cell division, macrophage senescence may contribute to the immune dysfunction observed in response to PM exposure.

The consequences of senescence vary considerably according to the initiating event and affected cell type. Manifestations of senescence include cell-cycle arrest, lysosome dysfunction, accumulation of oxidative stress, and the development of DNA damage (Childs et al., 2015). The pathogenic potential of senescent macrophages is highlighted in a recent report that the depletion of these cells improves outcomes in a KRAS-driven lung cancer model (Haston et al., 2023). Beyond the lungs, senescent macrophages have also been implicated in muscular dystrophy (Liu et al., 2022) and age-related adipose tissue dysfunction (Sawaki et al., 2023). The phenotypic heterogeneity of senescent cells necessitates the concurrent evaluation of multiple hallmarks of senescence that include cell cycle arrest, enhanced lysosomal mass and activity, lipid accumulation, and an aberrant secretory phenotype (González Gualda et al., 2021).

Here, we sought to evaluate the possibility that senescence is an outcome of uPM exposure in macrophages. While diesel exhaust particle (DEP) is often used as a surrogate for PM (Li, 2006), this may be an oversimplification as uPM is highly heterogeneous. Its composition varies according to city of origin and season (Kundu & Stone, 2014). Moreover, we have previously demonstrated that airway exposure of mice to uPM triggers a lung inflammatory response not observed in response to simpler particulates, like DEP or coal fly ash (Gour et al., 2018). To this end, we used PM collected from several urban locations and evaluated key metrics of cellular senescence. Unlike with sources of allergens (house dust mite [HDM] and ragweed), exposure of bone marrow-derived macrophages (BMDMs) to uPM results in the acquisition of a senescent profile. This offers insight into a potential mechanism by which uPM-mediated immune dysfunction may arise.

## Materials and methods

### Sources of particulate matter

Monosodium urate (MSU) and nano-sized silica oxide (nano-SiO_2_) were obtained from Invivogen. All PMs except for Baltimore were harvested using a sequential high-volume cyclone system that collected particles between 0.3-2.5 μm aerodynamic diameter when operated at a 1 m^3^/min flow rate, as described (Rule et al., 2010). Baltimore PM was collected with a modified high-volume cyclone system that collected particles between 0.3-10 μm aerodynamic diameter. DEP, also known as Standard Reference Material (SRM) 1650b, and urban particulate SRM 1648a were purchased from the National Institute for Standards and Technology (NIST). Coal fly ash (CFA) was purchased from Brandon Shores Unit power plant (Baltimore, MD). Particles were resuspended in PBS at 10 mg/mL. PM characteristics are summarized in Table 1.

**Table 1.** Characteristics of the particulate matter (PM) samples used in this study.

| **PM type**<br>**(source)** | **Collection system** | **Collection**<br>**date(s)** | **Size range** | **Physical / chemical Characterization** |
| --- | --- | --- | --- | --- |
| **SRM 1648a**<br>**(PM)** | Collected in St. Louis, MO in a purpose-built baghouse. Obtained from NIST | Collected from 1976-1977 | Bulk PM<br>Mean diameter = 5.85 µm | 13% carbon by mass, of which 10.5% is organic; high levels of metals (*e.g.* zinc, iron) and other inorganic elements. |
| **SRM 1650b**<br>**(DEP)** | Collected from four-cycle heavy duty diesel engines. Obtained from NIST | Collected in 1983 | Bulk PM<br>Mean diameter = 0.18 µm<br>Size between 0.12-0.33 µm | Derived from SRM1650a. |
| **Coal Fly Ash**<br>**(CFA)** | Brandon Shores Unit, Baltimore | Collected in 1998 | Size between 10-100 µm | The principal components of bituminous coal fly ash are silica, alumina, iron oxide with varying amounts of carbon. |
| **Phoenix PM** | Collected in Maricopa Co, AZ using a high-volume sequential cyclone* | Collected Jun – Jul 2008 | 0.3<d<2.5<br>Size between 0.3 and 2.5 µm | Characterized for metals and ions.<br>Mean concentration = 9.33±1.96 mg/m<sup>3</sup> |
| **Houston PM** | Collected in Harris Co, TX using a high-volume sequential | Collected Jan – Feb 2009 | 0.3<d<2.5<br>Size between 0.3 and 2.5 µm | Characterized for metals and ions.<br>Mean concentration = 8.68±2.98 |

|  | cyclone* |  |  | mg/m<sup>3</sup> |
| --- | --- | --- | --- | --- |
| **Pittsburgh PM** | Collected in Allegheny Co, PA using a high-volume sequential cyclone* | Collected Oct – Nov 2009 | 0.3<d<2.5<br>Size between 0.3 and 2.5 µm | Characterized for metals and ions.<br>Mean concentration = 10.32±2.81 mg/m<sup>3</sup> |
| **Baltimore PM<sub>10</sub>** | Collected in Baltimore, MD with a modified high-volume sequential cyclone<sup>+</sup> | Collected Fall 2012 | 0.3<d<10<br>Size between 0.3 and 10 µm | Characterized for metals and ions<br>C<br>Mean concentration = 14.3±7.5 mg/m<sup>3</sup> |

\* PM collected with sequential cyclone as described (Rule et al., 2010).
<sup>+</sup> Modified by eliminating the middle cyclone that would have collected PM between 2.5-10 µm

### Mice

C57Bl/6, *Nlrp3*<u>^-/-^</u>, *Casp1/4^-/-^*, and *Cd36*^-/-^ mice were obtained from Jackson Labs. All procedures were approved by the Animal Care and Use Committee of Johns Hopkins University.

### BMDM culture

Tibias and femurs from male C57Bl/6J mice were removed, and marrow harvested. Bone marrow cells were cultured on non-tissue culture-treated dishes and incubated in complete RPMI (cRPMI; 10% fetal bovine serum, penicillin/streptomycin, L-glutamine, and β-mercaptoethanol) supplemented with 15% L929-conditioned media for 7 days. BMDMs were washed once with PBS and harvested using Cell Stripper (Corning). For ELISAs, BMDMs were plated at 5×10^4^ cells/well of 96-well flat-bottom dish and left to adhere overnight in cRPMI. The following day, cells were exposed to particulates for 24h, and supernatants were harvested for ELISAs. For other assays, cells were seeded at 2×10^6^ cells/dish in small non-tissue culture-treated dishes and left to adhere overnight in cRPMI. Cells were then exposed to particulates for 24h and analyzed by flow cytometry.

### Chemicals

All inhibitors were purchased from Cayman Chemicals or Sigma.

### ELISA

Mouse IL-1α and TNFα were detected using DuoSets (RnD Systems).

### Flow cytometry

BMDMs were stained for viability using the Zombie Aqua Fixable dye (BioLegend), followed by Fc blocking using 20 µg/mL anti-CD16/32 (BioxCell, clone 2.4G2) for 20 min. Cells were stained with BV421-conjugated anti-CD64 (BioLegend, clone X54-5/7.1) and acquired on a BD LSRII. Data were analyzed using FlowJo v10 (BD Biosciences).

### C12FDG-based detection of ***β***-galactosidase activity

BMDMs were exposed to 100 nM bafilomycin A (Cayman Chemicals), then treated with 1 µM C12FDG (Cayman Chemicals) for 20 min at 37°C. Cells were washed and stained for surface markers, then analyzed by flow cytometry.

### Cell proliferation measured using the CellTiter-Glo assay

BMDMs were seeded at 1.0×10^4^ cells/well in a flat-bottom 96-well plate in cRPMI supplemented with 15% L929-conditioned media and stimulated as indicated. The CellTiter-Glo 2.0 assay (Promega) was then used to quantify ATP as a proxy for cell viability at 24h, 48h, and 72h post-stimulation.

### Statistical analysis

Data were analyzed via one-way analysis of variance (ANOVA) followed by Dunnett’s post-hoc test for multiple comparisons. Significance was defined as p<0.05. Analyses were executed using GraphPad Prism 9.

## Results

### Macrophages exposed to uPM acquire a senescent profile

We first wanted to determine whether uPM could drive manifestations of senescence in macrophages. To generate macrophages, we cultured bone marrow in differentiation media, yielding a pure macrophage culture, as determined by flow cytometry (Fig. 1A). One of the major determinants of this phenotype is the accumulation of senescence-associated β- galactosidase (SA-β-gal), a consequence of increased lysosomal mass. Cells were pre-treated with the vacuolar ATPase BafA1 to increase lysosomal pH, as SA-β-gal is often distinguished by its ability to function at a pH of up to 6 (*vs*. 4 for “normal” β-gal, though there is evidence of variation between cell types for β-gal activity (Noppe et al., 2009)). To assess for SA-β-gal activity, we used C12FDG, a fluorescent substrate for β-gal. We found that PMs from various urban sources and nano-SiO_2_ significantly increased SA-β-gal activity (Fig. 1B,C). This effect was not observed for particulates like DEP or CFA, or but only mildly for other common environmental exposures like house dust mite (HDM).

**Figure 1.**
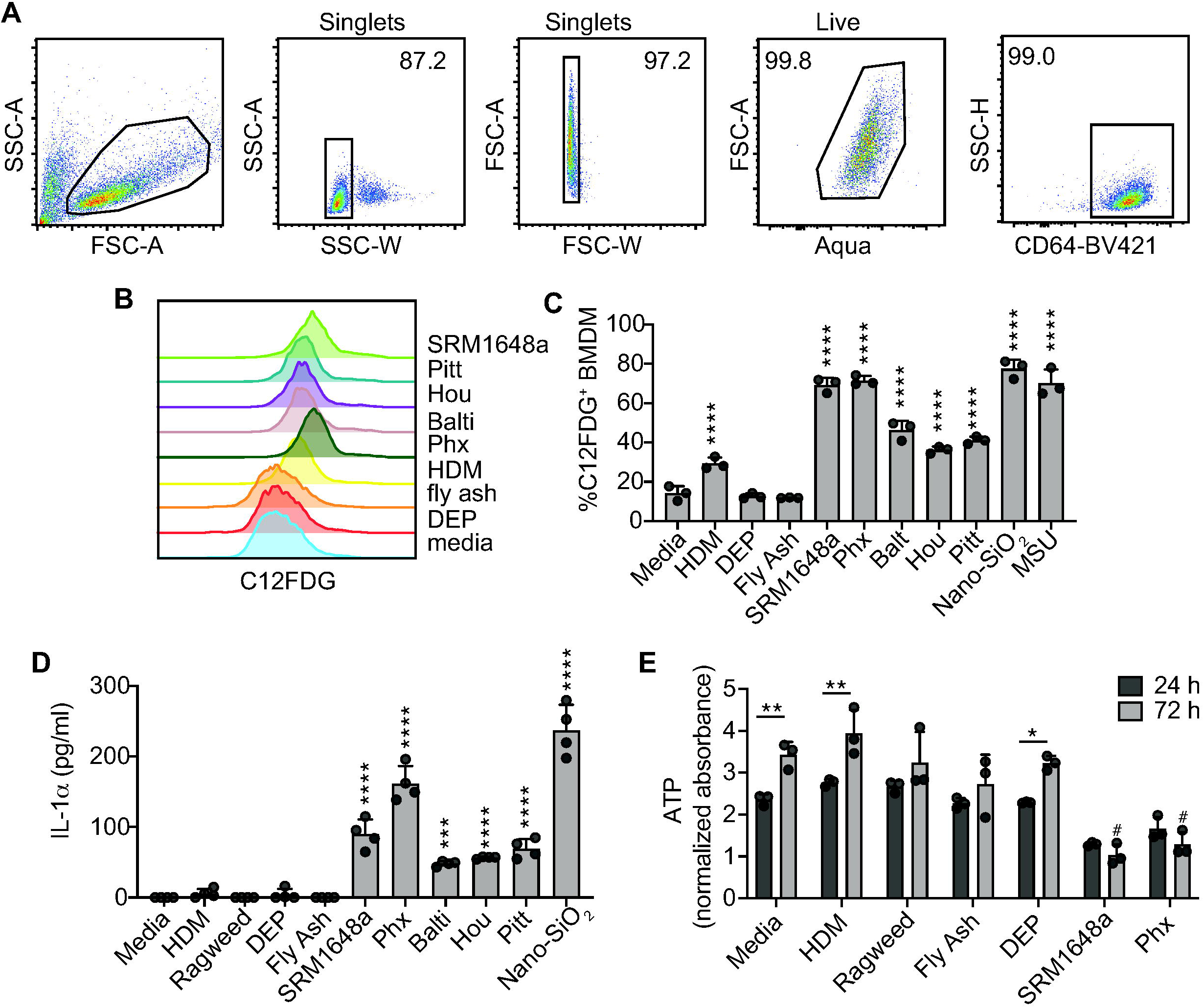
uPM drives senescence in macrophages. BMDMs were treated as indicated with media, sources of allergens (house dust mite [HDM] and ragweed), or particulates (diesel exhaust particulate [DEP], coal fly ash [CFA], SRM 1648a, Phoenix [Phx], Baltimore [Balti], Houston [Hou], Pittsburgh [Pitt], nanometer-sized silica oxide [nano-SiO_2_] or monosodium urate crystals [MSU]). Allergens were used at 50 µg/ml (based on protein content) and particulates at 100 µg/ml. (a) FlowJo gating schema, with CD64^+^ cells representing live BMDM singlets. (b) Frequency of C12FDG^+^ BMDMs 24h post-stimulation. (c) Proliferation was measured using intracellular ATP at 24h and 72h post-stimulation, normalized to baseline absorbance (0h). (d) Supernatant levels of IL-1α were quantified by ELISA 24h post-stimulation. Data plotted are technical replicates and are representative of at least two independent experiments. Data were analyzed using one-way ANOVA followed by Dunnett’s test for multiple comparisons. ****p<0.0001 versus media control.

In addition to SA-β-gal, senescent cells can adopt a specific secretome called the senescence-associated secreted phenotype (SASP). The SASP is characterized by the secretion of specific factors, notably the cytokine IL-1α (Coppé et al., 2010). IL-1α production is a salient characteristic of particulate exposure in macrophages and is a driver of particulate-induced inflammation (Rabolli et al., 2014). BMDMs were exposed to various particulates, including nano-SiO_2_, DEP and uPM, and two common sources of allergens, HDM and ragweed. Consistent with what we found for SA-β-gal, IL-1α secretion was induced by uPMs and nano-SiO_2_ but not by DEP, CFA, or allergens (Fig. 1D).

Finally, because senescent cells no longer divide, cell cycle arrest is another key manifestation of senescence (González Gualda et al., 2021). Quantification of ATP levels was used as a proxy for cell proliferation. BMDMs exposed to allergen (HDM and ragweed) displayed unimpeded proliferation over the 72-hour period (Fig. 1E). Conversely, uPM-treated cells showed a significant decrease in proliferation as compared to media control (Fig. 1E). These data suggest that exposure to uPM, but not allergens, promotes macrophage senescence.

### Senolytics abrogate senescence in uPM-treated BMDMs

Senolytics are a drug class identified for their ability to eliminate senescent cells, these include the FDA-approved tyrosine kinase inhibitor dasatinib (D) and the flavonoid quercetin (Q) (Kirkland & Tchkonia, 2020). We found that DQ-treated BMDM showed a significant reduction in uPM-induced SA-β-gal activity (Fig. 2A-C), as measured by frequency of SA-β-gal^+^ BMDMs (Fig. 2B) and median fluorescence intensity (MFI) of SA-β-gal in BMDMs (Fig. 2C). DQ also abrogated uPM-induced IL-1α (Fig. 2D) but had only a partial effect on uPM-induced TNFα (Fig. 2E). Reversal of uPM-induced SA-β-gal and SASP in response to DQ further supports the observation that exposure to urban particulates drive senescence.

**Figure 2.**
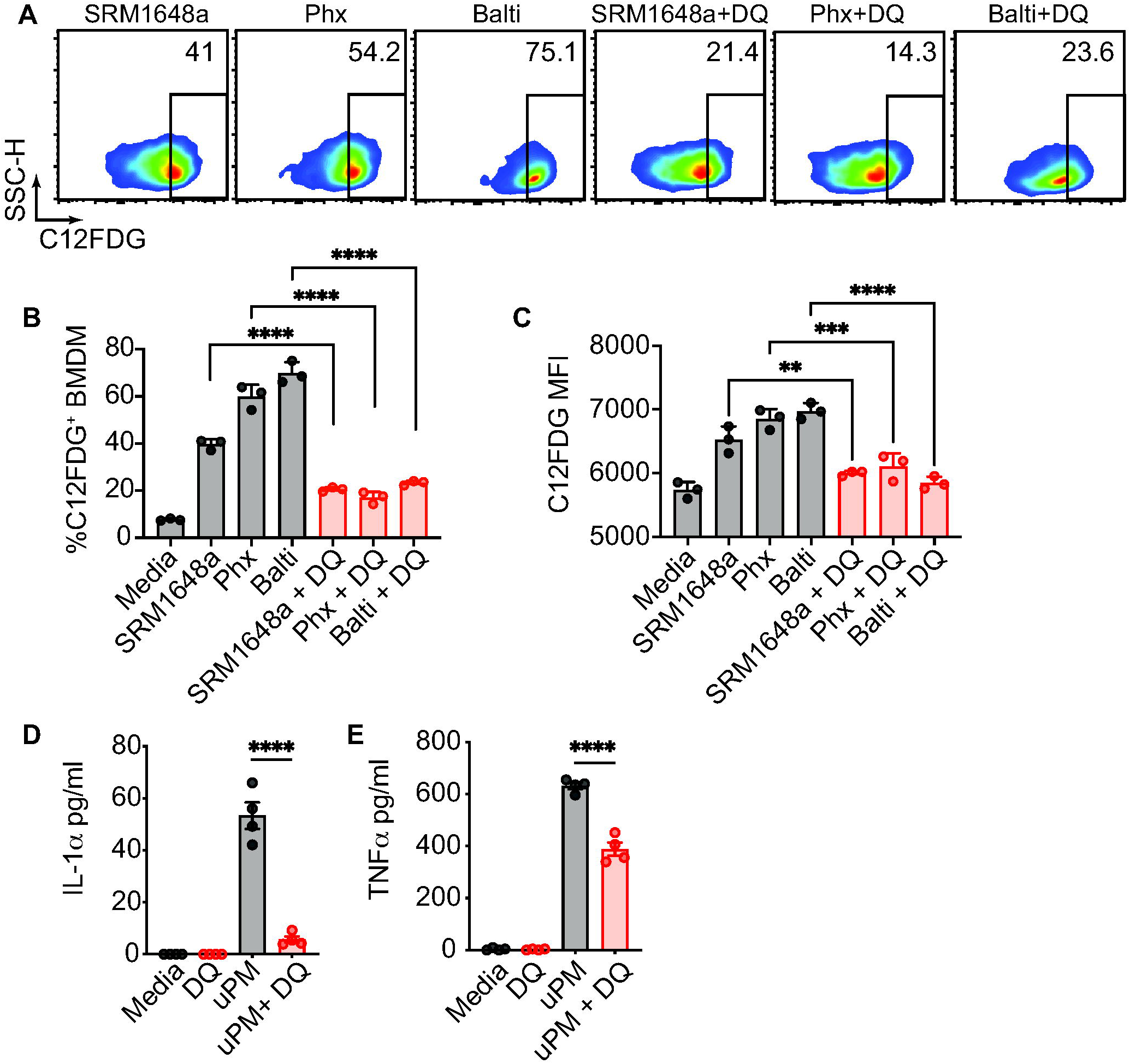
Senolytics abrogate uPM-induced senescence. BMDMs were treated with vehicle control or a cocktail of dasatinib (0.3 µM) and quercetin (30 µM) in combination with 100 µg/ml urban particulates as indicated. (a) Representative flow plots showing frequency of C12FDG^+^ cells 24h post-simulation, with corresponding (b) frequency of C12FDG+ cells and (c) median fluorescent intensity of C12FDG. Supernatant levels of (d) IL-1α and (e) TNFα were quantified by ELISA after stimulation with SRM 1648a (100 µg/ml, 24h). Data plotted are technical replicates and are representative of at least two independent experiments. Data were analyzed using one-way ANOVA followed by Dunnett’s test for multiple comparisons. ****p<0.0001.

### uPM-induced IL-1***α*** release is independent of the inflammasome, scavenger receptors, and NADPH oxidase-derived ROS

The inflammasome has been linked to the response of macrophages to PM (Sayan & Mossman, 2015). Thus, we wanted to ascertain the role of this pathway in mediating the adoption of the SASP in response to uPM. BMDMs from *Casp1/4*^-/-^ and *Nlrp3^-/-^* mice were evaluated for IL-1α secretion to ascertain the relevance of the inflammasome pathway in PM-mediated senescence. All inflammasome activators are thought to induce both IL-1α and IL-1β. These activators can be categorized as non-particulate (LPS, ATP, nigericin) or particulate (monosodium urate [MSU], alum, nano-SiO_2_). While non-particulate activators depend entirely on the inflammasome and caspase-1 for IL-1α secretion (Groß et al., 2012; Tsuchiya et al., 2021), we report that uPM exposure drives IL-1α independently of NLRP3 and caspase-1 activation (Fig. 3A,B). This is consistent with a previous report showing that silica- and MSU-induced IL-1α release is partially independent of inflammasome signaling (Groß et al., 2012).

**Figure 3.**
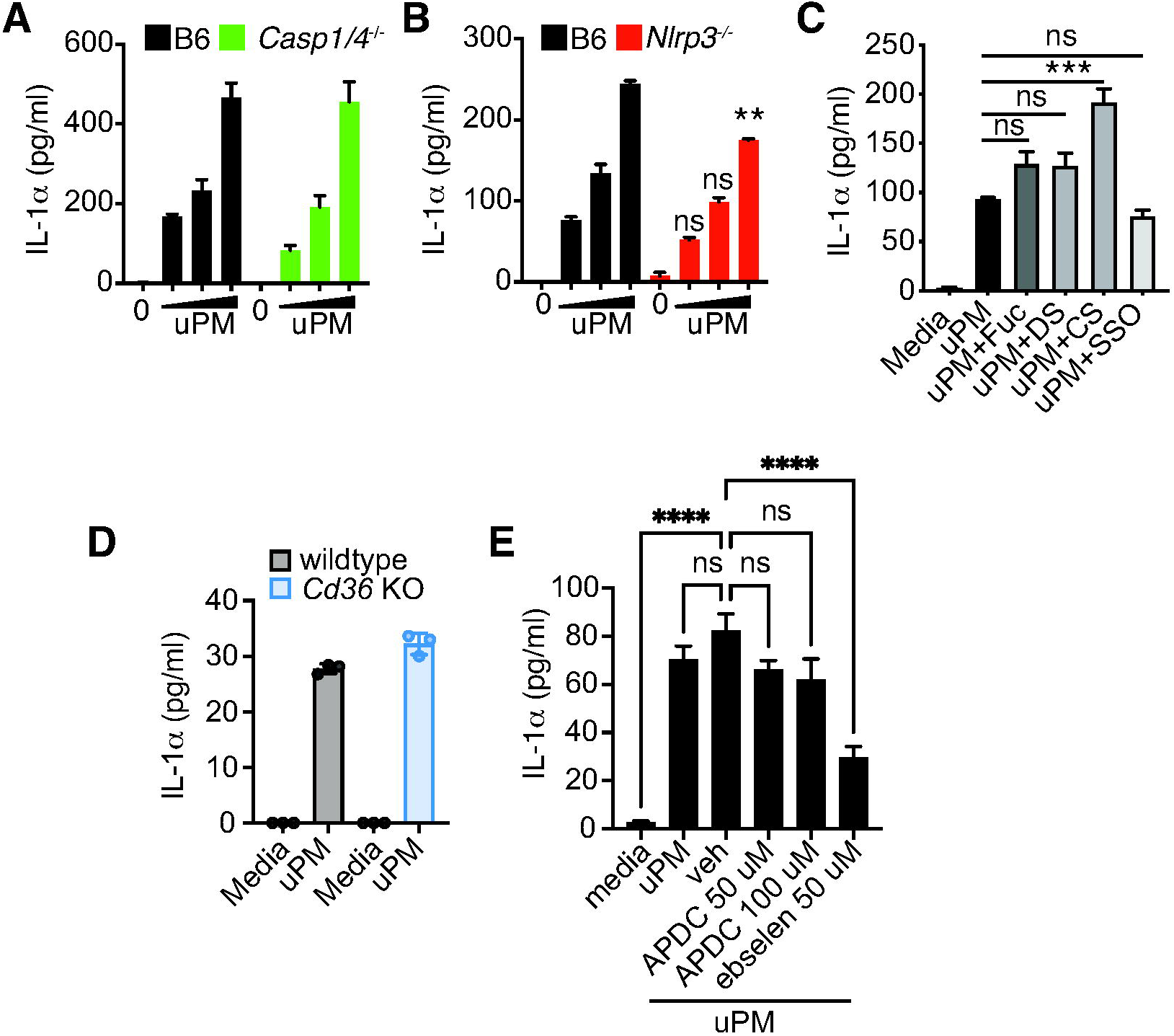
uPM-induced SASP is independent of caspase-1, NLPR3 and scavenger receptors. Levels of supernatant IL-1α after uPM stimulation (SRM 1648a, 100 µg/ml, 24h) of (a) B6 and *Casp1/4*^-/-^ BMDMs, and (b) B6 and *Nlrp3^-/-^* BMDMs. (c) Supernatant IL-1α levels from BMDMs pre-treated with scavenger receptor inhibitors before stimulation with uPM (SRM 1648a, 100 µg/ml, 24h). Scavenger receptor inhibitors fucoidan (Fuc, 100 µg/ml), dextran sulfate (DS, 100 µg/ml), and its control chondroitin sulfate (CS, 100 µg/ml) were used, as well as the CD36 inhibitor, sulfo-N-succidinimyl oleate (SSO, 200 µM). (d) Supernatant IL-1α levels from B6 and *Cd36*^-/-^ BMDMs stimulated with uPM (50 µg/ml). (E) Supernatant IL-1α levels from B6 BMDMs stimulated with uPM (50 µg/ml) in combination with vehicle (0.1% DMSO), APDC or ebselen. Data are plotted as mean+SEM and are representative of at least two independent experiments. Data were analyzed using one-way ANOVA followed by Dunnett’s test for multiple comparisons. ****p<0.0001.

Some groups have shown that scavenger receptors (SR) can interact with particulates, including silica, DEP, and titanium oxide (TiO_2_) particles (Arredouani et al., 2004; Hamilton et al., 2006; Levesque et al., 2013; Orr et al., 2011). While such particulates do not recapitulate the complexity of uPM, we wanted to test the contribution of SRs to the response observed on uPM exposure. We pre-treated BMDMs with broad inhibitors of SRs, which inhibit both class A and B receptors, then exposed cells to uPM. We found that IL-1α release was not dependent on SRs (Fig. 3C). Moreover, genetic knock-out of the SR, CD36, failed to block uPM-induced IL-1α production (Fig. 3D).

Reactive oxygen species (ROS) are an important marker of cellular stress and can potentiate the transition of a cell to a senescent phenotype (Davalli et al., 2016). To determine whether ROS plays a role in uPM-induced senescence, we treated BMDMs with the antioxidants ebselen and ammonium pyrrolidinedithiocarbamic acid (APDC). We found that blockade of NADPH oxidase-induced ROS via APDC did not prevent uPM-induced IL-1α release (Figure 3E). However, the blockade of peroxynitrite formation using ebselen partially reduced uPM-induced IL-1α. Overall, typical macrophage activation pathways do not regulate uPM-induced SASP.

### uPM-induced macrophage senescence requires intact phagolysosome function

We next examined other pathways that may regulate uPM-induced markers of senescence. As the formation of phagolysosomes is a critical step after phagocytosis, we sought to assess the importance of this pathway in uPM-induced senescence by using inhibitors targeting the phagolysosome pathway. First, the zinc chelator TPEN was used to disrupt lysosome function (Yu et al., 2019). Additionally, phagolysosome maturation was inhibited by blocking acidification of the luminal pH with BafA1, a vacuolar H^+^-ATPase inhibitor. Finally, the cathepsin B inhibitor CA-074Me was used to block proteolysis and undermine signaling pathways that maintain lysosome function and integrity (Xie et al., 2023). Blockade of these three pathways led to a reversal of uPM-induced SA-β-gal (Fig. 4A).

**Figure 4.**
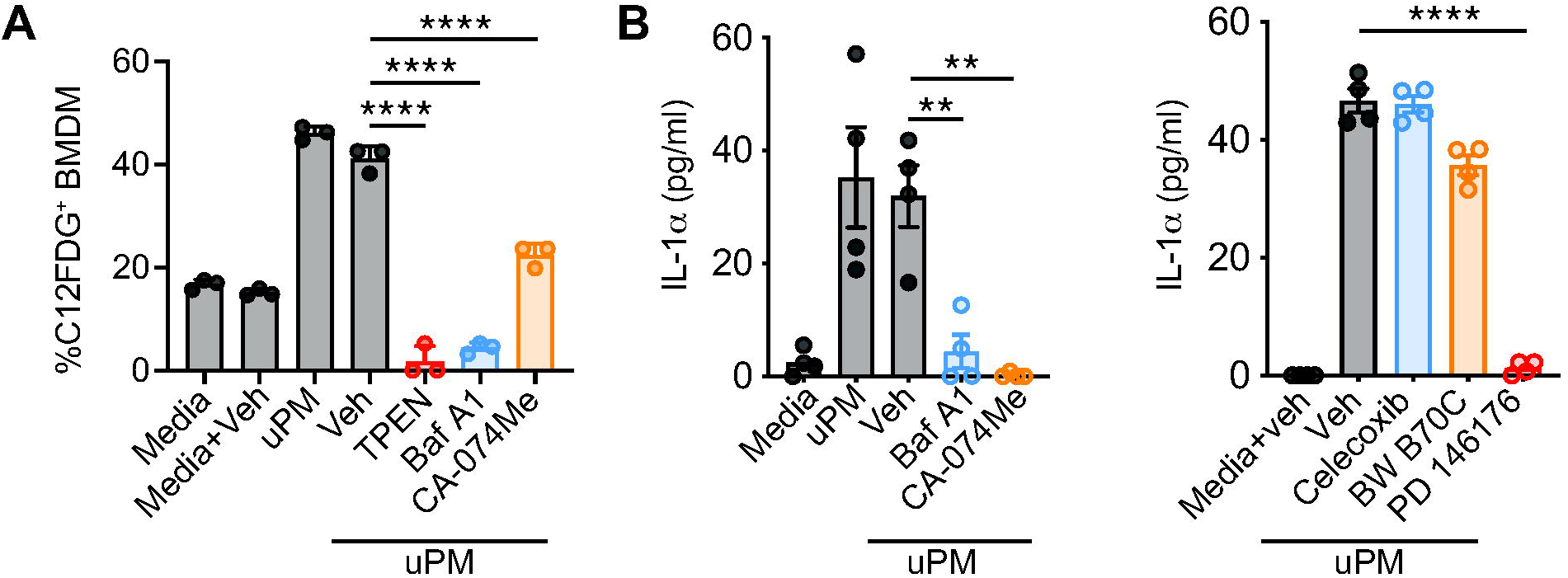
uPM-induced markers of senescence are dependent on a phagolysosome-15- lipoxygenase pathway. (a) Frequency of C12FDG^+^ BMDMs was analyzed in uPM-stimulated cells (100 µg/ml SRM 1648a, 24h) concurrently exposed to phagolysosome inhibitors: zinc chelator TPEN (20 µM), vacuolar ATPase inhibitor Bafilomycin A1 (BafA1, 40 nM), and cathepsin B inhibitor CA-074Me (10 µM). (b) Supernatant IL-1α levels from BMDMs stimulated with uPM (100 µg/ml SRM 1648a, 24h) and phagolysosome inhibitors BafA1 (40 nM) and CA-074-Me (10 µM). (c) Supernatant IL-1α levels from BMDMs stimulated with uPM (100 µg/ml SRM 1648a, 24h) and inhibitors of lipid pathway enzymes celecoxib (10 µM), BW B70C (10 µM), and PD 176146 (10 µM).

We next examined whether an intact phagolysosome pathway also affected uPM-induced SASP. BafA1-mediated lysosome basification and cathepsin B inhibition completely inhibited uPM- elicited IL-1α. (Fig. 4B). Lysosomes also have a role in lipid metabolism, and senescent cells often accumulate fatty acids, a substrate for enzymes including lipoxygenases (LO) and cyclooxygenases (COX) (Wiley & Campisi, 2016). To investigate this pathway, we treated cells with the 5-LO inhibitor, BW B70C, the 15-LO inhibitor PD 146176, and the COX-2 inhibitor, celecoxib. We found that inhibition of 15-LO, but not 5-LO or COX-2, ablated uPM-induced IL- 1α production (Fig. 4C).

Collectively, we propose that phagolysosome processing of uPM serves as a triggering event for macrophage senescence, as measured by SASP adoption, SA-β-gal activity, and cell cycle arrest (Fig. 5). Moreover, the mechanistic basis for uPM-induced senescence is independent of the inflammasome.

**Figure 5.**
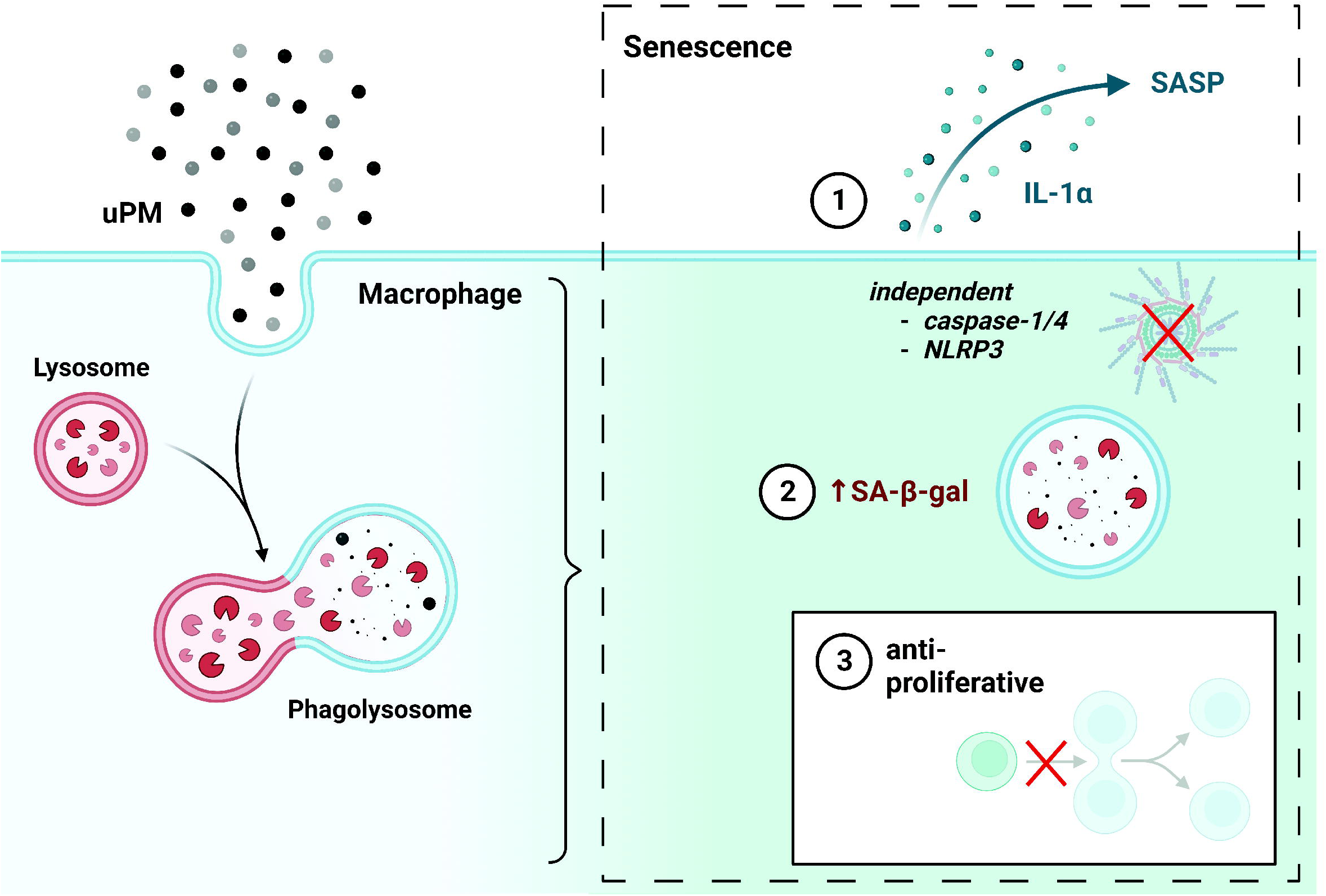
Phagolysosome processing of uPM triggers macrophage senescence. Macrophage phagocytosis and subsequent processing of uPM results in the acquisition of a cell profile indicative of senescence, as measured by SASP production (IL-1α), increased SA-β-gal activity, and inhibition of proliferation. This phenotypic shift is independent from caspase-1/4 and NLRP3. Image generated using Biorender.

## Discussion

Air pollution remains a significant public health challenge in both scope and severity, adversely affecting millions worldwide. Despite investigation into the health implications of PM exposure being in its nascent stages temporally, PM has been associated with myriad consequences to human health spanning diverse systems, including the immune system (Li et al., 2022; Thangavel et al., 2022; Ural et al., 2022). While PM has been proposed as a driver of dysregulated immunity, the mechanisms that give rise to dysfunction remain to be fully elucidated.

As critical sentinels in the lung, macrophages are among the first cell types to encounter airborne insults and are critical in coordinating the subsequent immune response (Franken et al., 2016; Ross et al., 2021). PM-induced senescence has been reported in the context of some cell types, including fibroblasts and epithelial cells (Jin et al., 2023; Sachdeva et al., 2019), but not macrophages. Our data demonstrates that exposure of macrophages to uPM results in senescence. These changes are uniquely observed in response to uPM and not other commonly used sources of PM (like DEP or CFA) or other environmental triggers like allergens.

Senolytics are a recent pharmacological tool for selective targeting of senescent cells. Murine studies into the therapeutic potential of DQ have demonstrated a reduction in senescent cells and improved functionality in myriad contexts, including diet-induced obesity, renal fibrosis, and osteoporosis (Chaib et al., 2022). In line with these data, we found that the manifestations of senescence induced in macrophages by uPM could be reversed by senolytics.

The relationship between inflammation and senescence is complex and likely bidirectional (Li et al., 2023). Blocking canonical pathways of macrophage activation, like the inflammasome, caspase-1/4, scavenger receptors, and ROS, failed to prevent uPM induction of SASP. This is in contrast to a report that caspase-4 drives LPS-induced SA-β-gal and cell cycle arrest in human fibroblasts (Fernández-Duran et al., 2022), supporting a distinction between the mechanisms by which classical inflammatory triggers (*e.g.,* LPS) and uPM drive senescence. Moreover, we found that phagolysosome activity drives uPM-induced senescence, suggesting phagolysosome processing of uPM by macrophages is key to the development of cellular senescence.

Senescence is associated with cytosolic accumulation of fatty acids, which are released by the lysosome and metabolized by various enzymes, including COX-2, 5-LO, and 15-LO. 15-LO is a known tumor suppressor linked to cellular senescence in prostate epithelial cells, as measured by SA-β-gal induction and cell cycle arrest (Suraneni et al., 2010). In line with this, the blockade of 15-LO function abrogated uPM-induced SASP.

Exposure to uPM may diminish macrophage functionality by driving senescence, thereby contributing to impaired tissue function. Our data warrant further exploration of uPM-induced macrophage senescence *in vivo*. Additionally, investigation of the functional ramifications of macrophage senescence is indicated to determine the mechanism(s) by which this phenomenon may contribute to disease.

## Author contribution

SAT performed the experiments with help from HMY and NG. AMR provided uPM samples. SAT, NG and SL designed the experiments and wrote the manuscript.

## Acknowledgements

This work was supported by the FHB33CRF- Maryland State Cigarette Restitution Fund (S.L.). S.L. also reports support from NIH R01AI27644 and R01AI170709 grants and the Johns Hopkins Catalyst Award.

